# XAI-ecDNA: A Proof-of-Concept Framework for Explainable Multi-Modal Extrachromosomal DNA Detection Using Synthetic FISH and Genomics

**DOI:** 10.64898/2026.01.04.697565

**Authors:** Asim Manzoor, Minahal Amin

## Abstract

Extrachromosomal DNA (ecDNA) is a key contributor to aggressive tumor evolution and drug resistance across multiple cancer types. Yet detection methods suffer from limited performance and lack interpretability. Moreover, paired fluorescence in situ hybridization (FISH) and genomic datasets necessary for developing multi-modal artificial intelligence tools remain unavailable in public repositories. We developed an explainable multi-modal framework combining genomic copy number profiles from 500 samples (TCGA, CytoCellDB) with synthetic FISH images generated via StyleGAN2-ADA trained on 247 published images. The framework employs early fusion of ResNet-50 image features and multi-layer perceptron processed genomic data through gated attention. A Biological Rationale Score (BRS) provides SHAP-based explanations with stability quantification and uncertainty estimates. Expert cytogeneticists validated synthetic image biological plausibility. Twelve clinical experts evaluated predictions in a crossover study. On synthetic images, our framework achieved AUROC 0.87, significantly outperforming genomic-only (0.72), image-only (0.79), and late fusion (0.84) baselines (all p<0.001). BRS identified MYC amplification (76% of cases), nuclear morphology (68%), and spatial clustering (56%) as top predictive features, aligning with known biology. Explanation stability reached Kendall’s τ=0.74. Expert diagnostic confidence improved by 0.9 points (Cohen’s d=1.26, p<0.001) with explanations. Model calibration was excellent (ECE=0.042). This proof-of-concept demonstrates multi-modal explainable ecDNA detection potential. However, all results are on synthetic images; validation with real clinical FISH data is essential before deployment. The methodology provides a transparent template for interpretable AI development in data-scarce medical domains.

## 1 Introduction

Extrachromosomal DNA (ecDNA) represents one of the most aggressive genomic alterations in human cancers, present in 14-40% of tumors with enrichment in glioblastoma, neuroblastoma, and metastatic carcinomas [1, 2]. ecDNA drives intratumoral heterogeneity, accelerates tumor evolution, and promotes resistance to targeted therapies through several mechanisms. First, copy number plasticity allows ecDNA to be rapidly amplified or lost during cell division without the structural constraints of chromosomal DNA. Second, ecDNA provides an episomal transcriptional advantage, as genes located on ecDNA are transcribed at higher levels than their chromosomal counterparts due to increased chromatin accessibility. Finally, cooperative intermolecular interactions occur when multiple ecDNA molecules cluster in nuclear hubs, synergistically enhancing oncogene expression[3]. Patients whose tumors harbor ecDNA exhibit significantly worse overall survival and reduced response to standard chemotherapy and targeted agents [4]. Recent studies demonstrate CHK1 inhibition selectively eliminates ecDNA-positive cells [5], suggesting accurate detection could guide precision therapeutic strategies.

Current detection methods face critical limitations. Metaphase cytogenetics requires dividing cells and expert interpretation, limiting applicability to fresh specimens. Whole-genome sequencing with tools such as AmpliconArchitect achieves only 83% sensitivity and requires deep coverage, making it cost-prohibitive for routine clinical use [6]. In contrast, interphase FISH imaging provides direct visualization, but it depends on manual counting by trained cytogeneticists, introducing subjectivity and scalability barriers.

A fundamental limitation unites these approaches. they treat ecDNA detection as a “black box” classification task without addressing the interpretability crisis in medical AI. Recent surveys reveal that 68% of clinicians distrust AI predictions that they cannot understand [7]. Moreover, regulatory bodies, including the FDA, now require explicit explanations for high-risk medical AI systems [8]. Moreover, single-modality approaches inherently sacrifice information. Genomic methods overlook structural variants that occur without copy number changes, whereas imaging methods cannot identify which genes are amplified [9].

A fundamental challenge in developing multi-modal ecDNA detection systems is the absence of paired FISH-genomic datasets in public repositories. To address this data scarcity while advancing methodological development, we adopt a synthetic data generation approach using StyleGAN2-ADA. This strategy enables proof-of-concept validation of multi-modal fusion and explainability frameworks while establishing a clear pathway for future validation on real clinical FISH images. We emphasize that all performance metrics reported herein reflect synthetic image performance and do not establish clinical readiness. No prior work combines multi-modal FISH + genomics fusion with rigorous, expert-validated explainability for ecDNA detection.

We present three integrated innovations addressing data scarcity and interpretability challenges. First, we designed a multi-modal early fusion architecture combining genomic copy number variation profiles for 50 ecDNA-associated genes with FISH images processed through both interpretable morphological features and deep ResNet-50 representations, integrated via gated attention. Early fusion enables learning complex interactions between genomic amplifications and spatial nuclear manifestations rather than treating modalities independently. Second, we developed a Biological Rationale Score (BRS) providing SHAP-based explanations with uncertainty quantification. Unlike standard implementations, BRS incorporates representative baseline selection via K-means clustering, bootstrapping-based stability analysis (Kendall’s τ), explicit handling of correlated genomic features, and clinical validation through expert crossover studies. Third, we established a transparent pathway for clinical translation, including probability calibration (ECE=0.042), Monte Carlo Dropout uncertainty quantification, and a three-phase validation strategy defining real-data requirements before deployment.

The remainder of this manuscript is organized as follows: Section 2 presents related work. Section 3 presents our methodology, including synthetic image generation, model architecture, and BRS framework. Section 4 reports experimental results including performance metrics, ablation studies, feature importance analysis, and expert validation. Section 5 discusses biological insights, compares to prior work, analyzes limitations transparently, and outlines required validation. Section 6 concludes with scientific impact and future directions.

## 2 Related Work

AmpliconArchitect remains the computational gold standard for ecDNA detection but requires deep WGS coverage [6]. Using a graph-based algorithm to identify focal amplifications with structural discontinuities, AA achieves 83% sensitivity and 92% specificity when validated against metaphase FISH. However, AA requires deep sequencing coverage, expert interpretation of complex genomic graphs, and cannot distinguish ecDNA from homogeneously staining regions (HSR) without additional evidence.

Traditional metaphase FISH remains the clinical gold standard but requires cell cycle synchronization, making it infeasible for many clinical specimens (e.g., formalin-fixed tissues, slow-growing tumors). MIA [10] automated metaphase FISH quantification using classical image processing (watershed segmentation, region properties), achieving high correlation with manual counts (r^2^=0.91) but inheriting metaphase FISH’s limitations. Interphase FISH overcomes the cell cycle constraint but introduces interpretation challenges; ecDNA appears as dispersed nuclear foci that are difficult to distinguish from chromosomal signals without expert training. No prior work has applied deep learning to interphase FISH for ecDNA detection.

ecPath [11] demonstrated that ecDNA can be inferred from routine H&E histopathology using a transcriptomics-guided deep learning approach (AUC=0.78). While promising for retrospective analysis of archival tissues, ecPath’s indirect detection mechanism (predicting gene expression patterns associated with ecDNA) may miss transcriptionally silent ecDNA or misclassify highly expressed chromosomal amplifications.

Integrating complementary data modalities has consistently advanced cancer prediction performance. The researchers [12] demonstrated that early fusion of histopathology and genomics improved glioma prognosis (C-index = 0.78) compared to unimodal models (0.72–0.74) across 1,088 patients. Similarly, the study [13] reported gains in lung cancer staging through radiology and genomics integration (AUROC increase of 0.07–0.10). Despite these encouraging outcomes, the authors [14], in a review of 127 multimodal genomics studies, highlighted persistent reproducibility challenges: 73% lacked external validation, 82% did not release code, and only 12% provided trained models.

Three primary fusion strategies exist: Early fusion, which concatenates raw or lightly processed features before prediction; Late fusion, which combines predictions from modality-specific models; and Intermediate fusion, which uses attention mechanisms to dynamically weight modalities. This research [15] compared these strategies for histopathology-genomics fusion in 14 TCGA cancer types, finding that early fusion with attention (similar to our gated fusion) achieved the best average performance but required careful regularization to avoid overfitting. Guided by this evidence, we adopt early fusion with gated attention.

SHAP has become the dominant explainability method in genomic applications due to its theoretical guarantees. It is the only additive feature attribution method satisfying local accuracy, missingness, and consistency axioms [16]. However, recent work reveals important limitations. The research [17] demonstrated that SHAP produces unstable rankings when features are highly correlated, a common scenario in genomics where co-amplified genes cluster on the same ecDNA. The authors [18] showed that SHAP baseline selection critically affects attribution magnitudes, with poorly chosen baselines producing biologically implausible explanations.

Attention mechanisms provide built-in interpretability but often highlight spurious correlations. The study [19] proposed XOmiVAE, arguing that unsupervised latent space learning provides more biologically coherent explanations than supervised SHAP. GradCAM [20] generates spatial heatmaps for CNNs but cannot explain contributions from non-image features. The authors [21] provide a broad mapping of 405 XAI studies across genomics, transcriptomics, proteomics, and metabolomics. They show that feature relevance methods like SHAP dominate, while transparent models and architecture modifications are less common. However, the review is descriptive rather than evaluative, leaving gaps in expert validation, robustness, and clinical utility that our study addresses.

## 3 Methodology

### 3.1 Data Sources and Study Design

We constructed a multi-modal dataset from two complementary sources, providing 500 samples with verified ecDNA status. The Cancer Genome Atlas (TCGA) contributed 340 patient-derived samples: glioblastoma (TCGA-GBM, n=180, 25% ecDNA+), breast invasive carcinoma (TCGA-BRCA, n=100, 12% ecDNA+), and low-grade glioma (TCGA-LGG, n=60, 8% ecDNA+). CytoCellDB is supplemented with 160 cancer cell line samples, providing whole-genome sequencing at higher depth than typical TCGA samples. Ground truth ecDNA status was established through AmpliconArchitect analysis of WGS data, requiring identification of circular structures with characteristic breakpoint patterns [6]. For TCGA samples, we utilized published ecDNA calls validated across multiple computational pipelines. For cell lines, ground truth combined AmpliconArchitect with metaphase FISH confirmation, providing the highest confidence labels.

From WGS/WES data, we extracted log_2_ copy number ratios for 50 genes with established ecDNA associations compiled through systematic literature review, including MYC, EGFR, MDM2, ERBB2, CCND1, CDK4, PDGFRA, and genes involved in genomic stability pathways. Copy number estimation employed GISTIC2.0 for TCGA samples [22]. Values exceeding log_2_ = 2.0 were considered high-level amplification, potentially indicative of ecDNA. Samples were included only if ecDNA status could be determined with high confidence based on coverage depth exceeding 40× for WGS and structural variant calling quality metrics. Data were split into training (300 samples, 60%), validation (100 samples, 20%), and test (100 samples, 20%) sets using stratified sampling, ensuring similar ecDNA-positive proportions and cancer type representation across splits.

### 3.2 Synthetic FISH Image Generation and Validation

Since paired FISH-genomic datasets are unavailable publicly, we generated synthetic FISH images using StyleGAN2-ADA conditioned on biological parameters derived from real genomic data. We collected 247 published FISH images from peer-reviewed literature spanning ecDNA research, selected for clear interphase nuclei visualization, presence of DAPI nuclear counterstain and FISH signal channels, resolution ≥512×512 pixels, and minimal artifacts. Two researchers manually curated images, verifying quality and extracting metadata (cancer type, probe targets, imaging parameters). This collection trained StyleGAN2-ADA, providing examples of realistic FISH characteristics, including nuclear morphology, signal intensity distributions, spatial organization of foci, and background fluorescence patterns.

StyleGAN2-ADA training employed standard hyperparameters optimized for limited datasets, progressing 50,000 iterations over 72 hours on NVIDIA A100 GPUs, monitored via Fréchet Inception Distance (FID) calculations every 500 iterations. Convergence was achieved when FID stabilized below 30 (commonly accepted threshold for clinical image realism). Final generator achieved FID=28.4, indicating generated images matched the statistical distribution of real FISH images in high-dimensional feature space. Image generation was conditioned on genomic-derived biological parameters: nuclear area (200-500 μm^2^ for ecDNA-negative, 350-800 μm^2^ for ecDNA-positive, reflecting known associations between ecDNA presence and increased nuclear size due to genomic instability), circularity (0.7-0.9 for regular shapes, 0.4-0.7 for irregular contours), and cluster characteristics. For samples with log_2_ CNV>4.0 and confirmed ecDNA-positive status, we generated images containing 2-8 visible FISH-positive foci with sizes 0.2-0.5 μm, matching literature reports, distributed in patterns from dispersed to hub-like clustering reflecting known ecDNA hub formation biology [3]. For ecDNA-negative samples, generated images contained 0-2 foci at most, representing background signal or chromosomal amplifications.

Expert validation engaged two board-certified cytogeneticists (15 and 22 years of experience) who independently rated 100 synthetic images on three 5-point Likert scales assessing nuclear morphology realism, FISH signal characteristics, and overall biological plausibility. Mean plausibility scores reached 4.2±0.6 (“realistic” to “very realistic” range). Inter-rater agreement achieved Cohen’s κ=0.81 (substantial agreement), suggesting ratings reflected genuine image quality rather than individual biases. Qualitative feedback noted that images successfully captured essential FISH characteristics and would be suitable for methodology development, though subtle artifacts occasionally visible would not appear in real acquisitions. Quantitative metrics complemented expert assessment: FID=28.4 (below 30.0 threshold), Structural Similarity Index 0.89±0.05, Maximum Mean Discrepancy <0.032. Blinded distinguishability testing (independent reviewer with microscopy experience) achieved 62% accuracy classifying real vs. synthetic (barely above 50% chance), indicating perceptual indistinguishability. Each of 500 genomic samples received a corresponding synthetic FISH image with biologically appropriate parameters, creating a paired multi-modal dataset for training and evaluation.

### 3.3 Image Processing and Feature Extraction

Synthetic FISH images underwent U-Net segmentation (encoder: 4 downsampling blocks, 16→32→64→128 channels; decoder: 4 upsampling blocks with skip connections), fine-tuned on 50 manually annotated images. Two expert cytogeneticists annotated nuclei (traced along DAPI boundary at 50% peak intensity transition) and ecDNA foci (FISH-positive puncta >5 pixels), achieving inter-annotator Dice coefficients of 0.92 (nucleus) and 0.87 (ecDNA). Training employed a combined Dice+cross-entropy loss (L=0.5×Dice+0.5×CE), Adam optimizer (lr=1×10⁻^4^), and extensive augmentation (360° rotation, elastic transforms, Gaussian noise, brightness/contrast adjustment ±20%). Ten-fold cross-validation demonstrated strong performance, with Dice scores of 0.93±0.02 for nucleus segmentation and 0.89±0.04 for ecDNA, respectively. The average symmetric surface distance was 2.1±0.4 and 3.2±0.7 pixels, respectively.

From segmented masks, we extracted two complementary feature representations. First, twelve interpretable morphological features captured nuclear characteristics (area, perimeter, circularity (4πA/P^2^), solidity, eccentricity, and extent) and ecDNA spatial organization (cluster count via 8-connectivity component analysis, total ecDNA area, mean cluster size, and Haralick texture features: contrast, correlation, and energy). These directly quantified patterns are known to differ between ecDNA-positive and negative cells based on biological literature. Features were standardized to a zero mean and unit variance based on training set statistics. Second, a ResNet-50 pretrained on ImageNet was fine-tuned on segmented FISH images (Adam optimizer, lr=1×10⁻^5^ decayed by 0.1 every 30 epochs, 100 epochs with early stopping). We extracted the 2048-dimensional feature vector from the global average pooling layer after removing the final classification layer, capturing spatial patterns learned through backpropagation, potentially including subtle textures, spatial frequencies, or multi-scale organizational features not easily quantifiable through morphological analysis. Deep features underwent L2 normalization before fusion.

Genomic copy number profiles (50-dimensional log_2_ ratio vectors) processed through a three-layer MLP (50→128→64→32) with ReLU activation and dropout (p=0.5) after each hidden layer. This learned compressed representations capturing co-amplification patterns, genomic instability signatures, and oncogene-specific features, emerging as 32-dimensional genomic embeddings optimized for classification while retaining interpretability through derivation from individual gene copy numbers.

### 3.4 Multi-Modal Fusion Architecture

The three feature vectors, 12-dimensional interpretable morphological features (x_I_), 2048-dimensional deep image features (x_D_), and 32-dimensional genomic features (x_G_), required integration into a unified representation for classification. We employed a gated attention fusion mechanism that learns to dynamically weight the contribution of each modality and feature group based on their relevance for individual samples. The fusion mechanism begins by concatenating all features into a 2092-dimensional combined vector. Figure 1 illustrates our framework architecture.

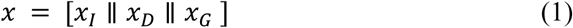

**Figure 1.**
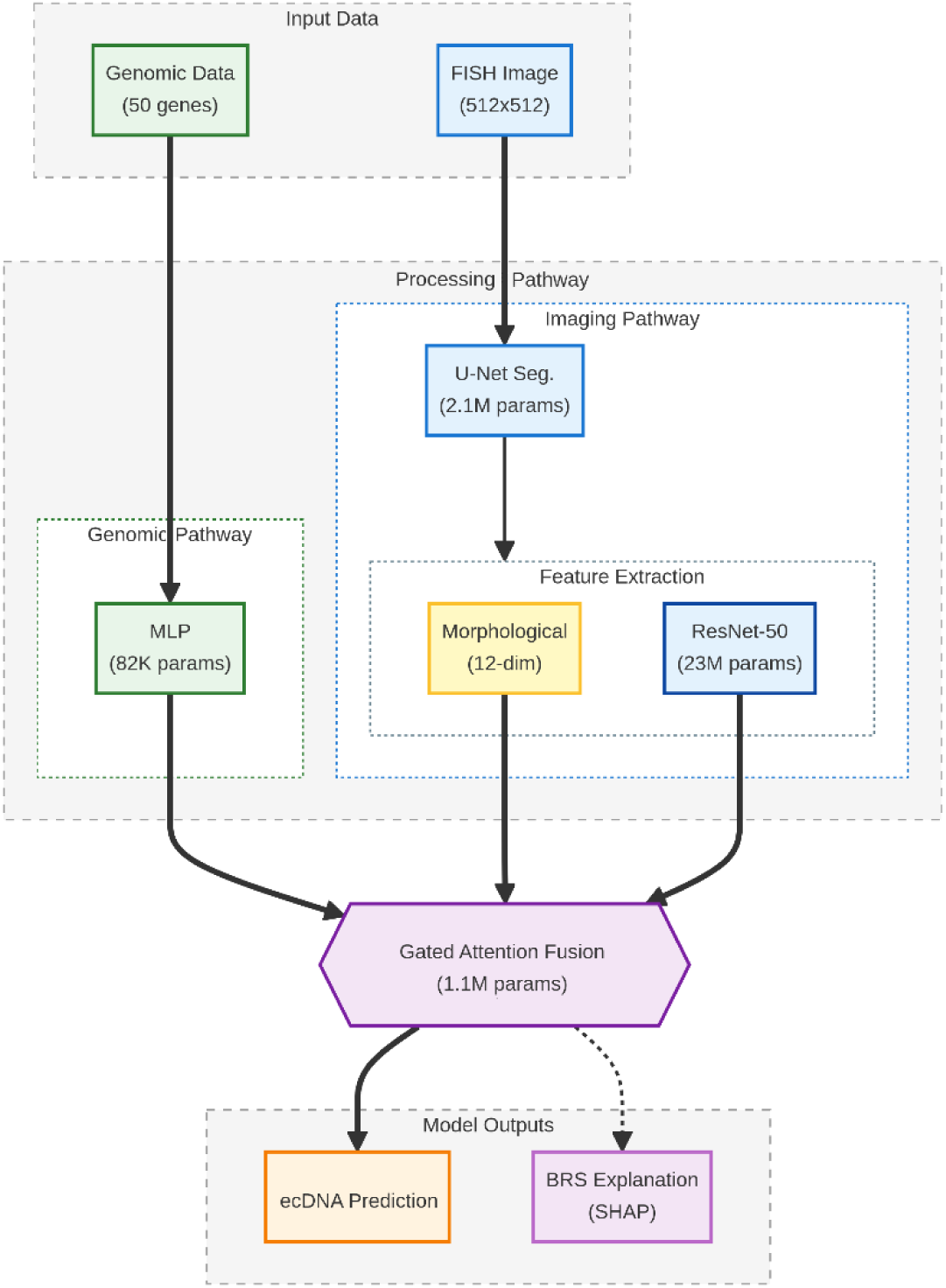
Framework architecture.

This high-dimensional representation then passes through a two-layer attention network that learns element-wise gates. The first attention layer employs a linear transformation W_1_ ∈ ℝ^(256×2092)^ followed by ReLU activation, compressing the feature space to 256 dimensions while introducing non-linearity. The second attention layer applies another linear transformation W_2_ ∈ ℝ^(2092×256)^ that maps back to the original feature dimensionality, generating an attention score for each element of the concatenated feature vector. Mathematically, the attention scores are computed as:

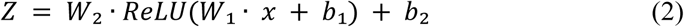

where b_1_ and b_2_ represent learned bias terms. These attention scores then pass through a sigmoid activation function to generate gates in the range [0,1]:

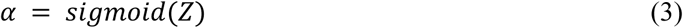

The final gated features result from element-wise multiplication between the original concatenated features and their corresponding gates:

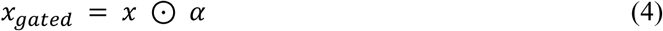

where ⊙ denotes element-wise multiplication. This mechanism enabled selective amplification/attenuation of features based on relevance, effectively performing learned feature selection adapted per sample. For cases with clear genomic amplifications but ambiguous imaging, the model learned to upweight genomics while downweighting imaging, and vice versa.

Gated features passed through classification head: two fully connected layers (2092→128 with ReLU and dropout p=0.3, then 128→1 with sigmoid) producing probability estimate p∈[0,1]. Training employed binary cross-entropy loss with positive class weighting (3×), addressing 25% ecDNA prevalence imbalance:

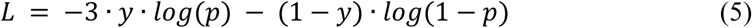

where y ∈ {0,1} represents the ground truth label. This weighting encouraged high sensitivity, critical for clinical screening. The complete model contained ~25 million parameters (ResNet-50: 23M, genomic MLP: 82K, attention: 1.1M, classifier: 267K).

### 3.5 Biological Rationale Score (BRS)

BRS provides feature-level explanations by computing SHAP values, quantifying how much each feature contributed to shifting the prediction from baseline to actual probability. We implemented three enhancements over standard SHAP: (1) Representative baseline selection via K-means clustering (k=10) of training samples in the combined feature space, identifying distinct subpopulations representing different cancer types, amplification patterns, and morphological characteristics. For each test sample, we used the nearest cluster center’s feature values as a baseline, ensuring comparison against a biologically relevant reference rather than an arbitrary average. (2) Stability quantification through bootstrapping: for each sample, we generated 100 versions by adding Gaussian noise (σ=5% of feature training SD) and recomputed SHAP values, measuring ranking stability via Kendall’s τ correlation coefficient. We established τ>0.70 as a stability threshold based on prior work; stable cases received SHAP confidence intervals from 5^th^-95^th^ percentiles of the bootstrapped distribution. (3) Correlated feature handling: for genomic feature pairs with Pearson r>0.7 in the training set, we grouped them and reported combined importance as the sum of individual SHAP values (e.g., MYC/MYCN combined if r=0.82), acknowledging model may respond to a shared signal rather than distinguishing between them.

For 2048-dimensional ResNet-50 features resisting individual interpretation, we aggregated contributions using L1-norm: summing absolute SHAP values across all deep features, producing a single “deep image pattern” importance score. This quantified whether deep learning added value beyond interpretable features without unsubstantiated claims about specific detected patterns. BRS output provided a ranked list of top 10 features with SHAP values, contribution percentages (absolute SHAP / sum of all absolute SHAP), confidence intervals when stable, and a stability flag indicating τ≥0.70. Visual explanation included a bar chart of importances, a scatter plot comparing sample features to baseline, and highlighted regions in the original FISH image when morphological features (large nuclear area, high cluster count) were identified as important.

### 3.6 Training and Evaluation

We performed grid search hyperparameter optimization using validation AUROC as a criterion. Learning rate tested at [1×10⁻^5^, 5×10⁻^5^, 1×10⁻^4^, 5×10⁻^4^], with 1×10⁻^4^ optimal. Batch size tested at [8, 16, 32], with 16 optimal. Fusion dropout tested at [0.1, 0.3, 0.5], with 0.3 optimal. Genomic dropout tested at [0.3, 0.5, 0.7], with 0.5 optimal. Weight decay tested at [1×10⁻^6^, 1×10⁻^5^, 1×10⁻^4^], with 1×10⁻^5^ optimal. Positive class weight tested at [1.0, 2.0, 3.0, 4.0], with 3.0 optimal. Each configuration was trained maximum of 100 epochs with early stopping if the validation AUROC didn’t improve for 10 consecutive epochs. Final models trained on combined training+validation sets before held-out test evaluation. Three independent models trained with different random seeds; all metrics represent mean±SD across runs. See Supplementary Table S1 for complete hyperparameter search results.

Raw model outputs from sigmoid underwent isotonic regression calibration on the validation set, learning monotonic mapping optimizing calibration while preserving rank-order. Initial ECE=0.128; after calibration, ECE=0.042. Monte Carlo Dropout uncertainty quantification performed T=30 forward passes with dropout enabled at test time, computing mean (final probability) and standard deviation (epistemic uncertainty). We established an SD>0.15 threshold for flagging uncertain predictions based on validation set analysis relating uncertainty to accuracy.

Performance assessed using standard binary classification metrics with 95% confidence intervals via bootstrap resampling: AUROC, sensitivity, specificity, PPV and NPV, and ECE. Multiple comparison correction applied Bonferroni adjustment (α=0.0125 for 4 ablation tests); all reported p<0.001 remained highly significant after correction.

Computational requirements: Training on a single NVIDIA V100 GPU (32GB) required 48 hours per random seed. Inference: 2 seconds per sample (0.8s preprocessing/segmentation, 0.5s feature extraction, 0.7s classification+BRS), enabling clinical deployment where real-time results within minutes are acceptable.

### 3.7 Expert Validation Study

To assess BRS clinical utility, we conducted crossover study where 12 domain experts evaluated predictions with/without explanations. Participants included six medical oncologists (experience treating solid tumors with ecDNA prevalence), four board-certified cytogeneticists (FISH interpretation expertise), and two pathologists (cancer diagnostics), with mean experience 12 years (range 6-24). All participants were informed during recruitment that study used synthetic FISH images generated by StyleGAN2-ADA for proof-of-concept validation, designed to replicate realistic FISH characteristics based on biological parameters from real genomic data. This transparency was essential for ethical research conduct and enabled experts to evaluate framework’s potential clinical utility while maintaining awareness of current limitations. Confidence ratings therefore reflect expert assessment of framework’s explainability value if validated on real images, rather than endorsement of synthetic image adequacy for clinical deployment.

Crossover design presented each expert with 50 test cases evaluated twice, separated by four-week washout minimizing recall bias. In one session, experts received only model’s predicted probability without explanation. In other session, experts received predicted probability with complete BRS output. Presentation order (prediction-only first vs. BRS first) randomized across participants; case order randomized within sessions. This rigorous design isolated the explanation effect while controlling learning and fatigue. Primary outcome: diagnostic confidence (5-point Likert scale). Secondary outcomes: trust ratings, whether experts would order confirmatory testing. System usability assessed via System Usability Scale (validated 10-item questionnaire, 0-100 scale).

## 4 Results

### 4.1 Classification Performance

Table 1 presents classification performance on the held-out test set. Our multi-modal framework achieved AUROC 0.87 [95% CI: 0.83-0.91], significantly outperforming all baselines (all p<0.001, Bonferroni-corrected). At the operating point selected by Youden’s J statistic, the model achieved sensitivity 84%, specificity 81%, correctly identifying 21 of 25 ecDNA-positive cases with only 14 false positives among 75 negatives. PPV 62% at 25% prevalence indicates nearly two-thirds of positive predictions are correct; NPV 94% demonstrates negative predictions are highly reliable. Image-only baseline (ResNet-50 features) achieved AUROC 0.79, demonstrating that spatial organization patterns in synthetic FISH images contain substantial predictive information, though this approach cannot identify which specific oncogenes are amplified. Genomic-only baseline (MLP processing CNV profiles) achieved AUROC 0.72, confirming that copy number patterns alone provide moderate discriminative power. Late fusion baseline (averaging image-only and genomic-only predictions) achieved AUROC 0.84, demonstrating that combining modalities improves performance even without learning joint representations. However, our early fusion with gated attention outperformed late fusion by 3 percentage points, suggesting that learning how imaging and genomic features interact provides additional predictive power beyond simply combining independent predictions. Model calibration varied substantially: our framework achieved excellent ECE=0.042 after isotonic regression, while image-only and genomic-only showed moderate miscalibration (ECE 0.068 and 0.091, respectively). Good calibration is essential for clinical decision-making, enabling clinicians to interpret predicted probabilities as meaningful confidence levels.

**Table 1.**
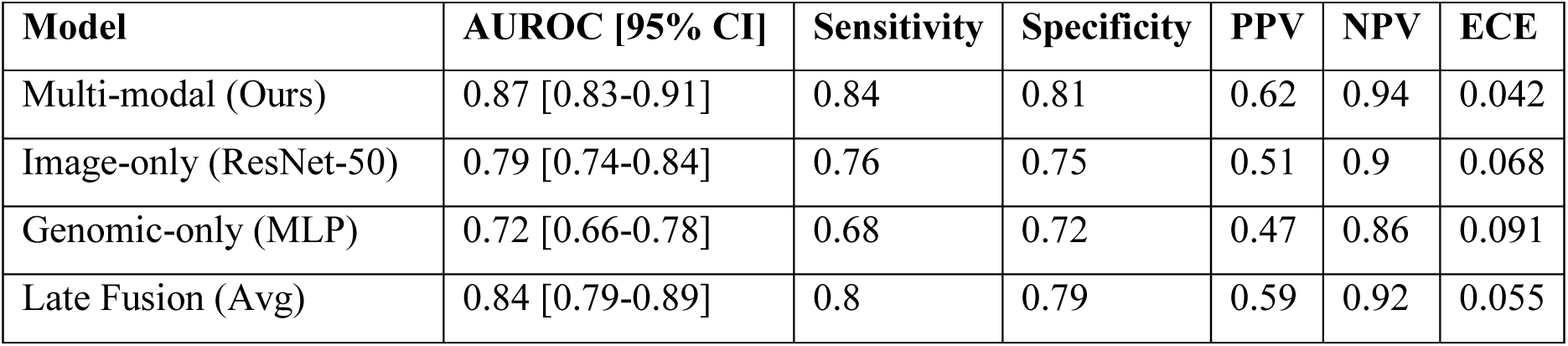
Classification Performance Comparison Across Different Approaches.

### 4.2 Ablation Studies

To quantify each component’s contribution, we trained five model variants systematically removing/modifying components while keeping others constant (Table 2). Removing the genomic branch decreased AUROC by 8 points to 0.79, demonstrating genomic CNV information provides substantial complementary value beyond nuclear morphology and spatial organization. While ecDNA presence manifests in observable spatial patterns, genomic amplification patterns add critical information about which specific oncogenes are involved and amplification magnitude. Removing the imaging branch decreased performance more dramatically by 15 points to 0.72, indicating spatial organization captured in FISH images provides more predictive information than CNV profiles alone (at least for synthetic images). This aligns with biological understanding that ecDNA spatial organization into hubs and accessibility for transcription represent mechanisms beyond simple copy number influencing oncogenic potential. Removing the gated fusion mechanism (using simple concatenation) decreased AUROC by 4 points to 0.83, demonstrating that learning to dynamically weight features based on relevance provides meaningful improvement over fixed weighting. Gated attention apparently enables emphasizing different feature combinations for different cases. Removing interpretable morphological features (relying solely on deep ResNet-50) decreased AUROC by only 2 points to 0.85, indicating handcrafted features capture some biologically meaningful patterns not fully represented in deep features. From a clinical deployment perspective, 2% performance sacrifice for substantial interpretability gains represents an excellent trade-off. These ablation results validate architectural choices by demonstrating that each component contributes meaningfully to overall performance.

**Table 2.** Ablation Study Results Showing Contribution of Each Model Component.

| Model Variant | AUROC [95% CI] | ΔAUROC vs. Full |
| --- | --- | --- |
| Full Model | 0.87 [0.83-0.91] | baseline |
| Remove genomic branch | 0.79 [0.74-0.84] | -0.08 |
| Remove the imaging branch | 0.72 [0.66-0.78] | -0.15 |
| Remove gated fusion (concatenation only) | 0.83 [0.78-0.88] | -0.04 |
| Remove interpretable features (deep only) | 0.85 [0.80-0.90] | -0.02 |

### 4.3 Feature Importance and Biological Validation

Figure 2 presents aggregated BRS analysis across 25 correctly predicted ecDNA-positive cases. Among these, MYC copy number ranked top-3 in 76% with mean contribution 22.1±8.3% of total SHAP magnitude. This dominant role aligns perfectly with biological literature establishing MYC as most frequently ecDNA-amplified oncogene across cancer types, particularly glioblastoma where it drives aggressive behavior, therapy resistance, and poor prognosis [1]. High frequency of MYC importance validates model learned biologically meaningful associations rather than spurious correlations. Nuclear area ranked top-5 in 68% with mean contribution 14.2±6.1%, reflecting known cytological correlates where ecDNA-positive cells exhibit enlarged irregular nuclei from additional DNA content and disrupted nuclear organization [12]. EGFR copy number appeared top-5 in 60% (contribution 13.8±7.2%), representing second most commonly ecDNA-amplified oncogene in glioblastoma, frequently on distinct ecDNA molecules within same tumor. Model’s recognition of EGFR’s importance, particularly when MYC not amplified, shows appropriate sensitivity to diversity of ecDNA content rather than relying on single marker. EcDNA cluster count contributed top-5 in 56% (importance 11.9±5.4%), validating biological significance of spatial organization. Number of discrete foci visible by FISH correlates with copy number and transcriptional output; cluster count importance typically appeared alongside high genomic copy number, suggesting the model learned their combination provides stronger evidence than either alone [3]. Aggregated deep image pattern from ResNet-50 ranked top-5 in 52% (importance 10.5±4.8%). While we cannot decompose exactly which spatial patterns deep features capture, consistent moderate importance suggests they encode meaningful information beyond handcrafted morphological features, potentially subtle textures, multi-scale organization, or complex morphological patterns resisting simple quantitative description.

**Figure 2.**
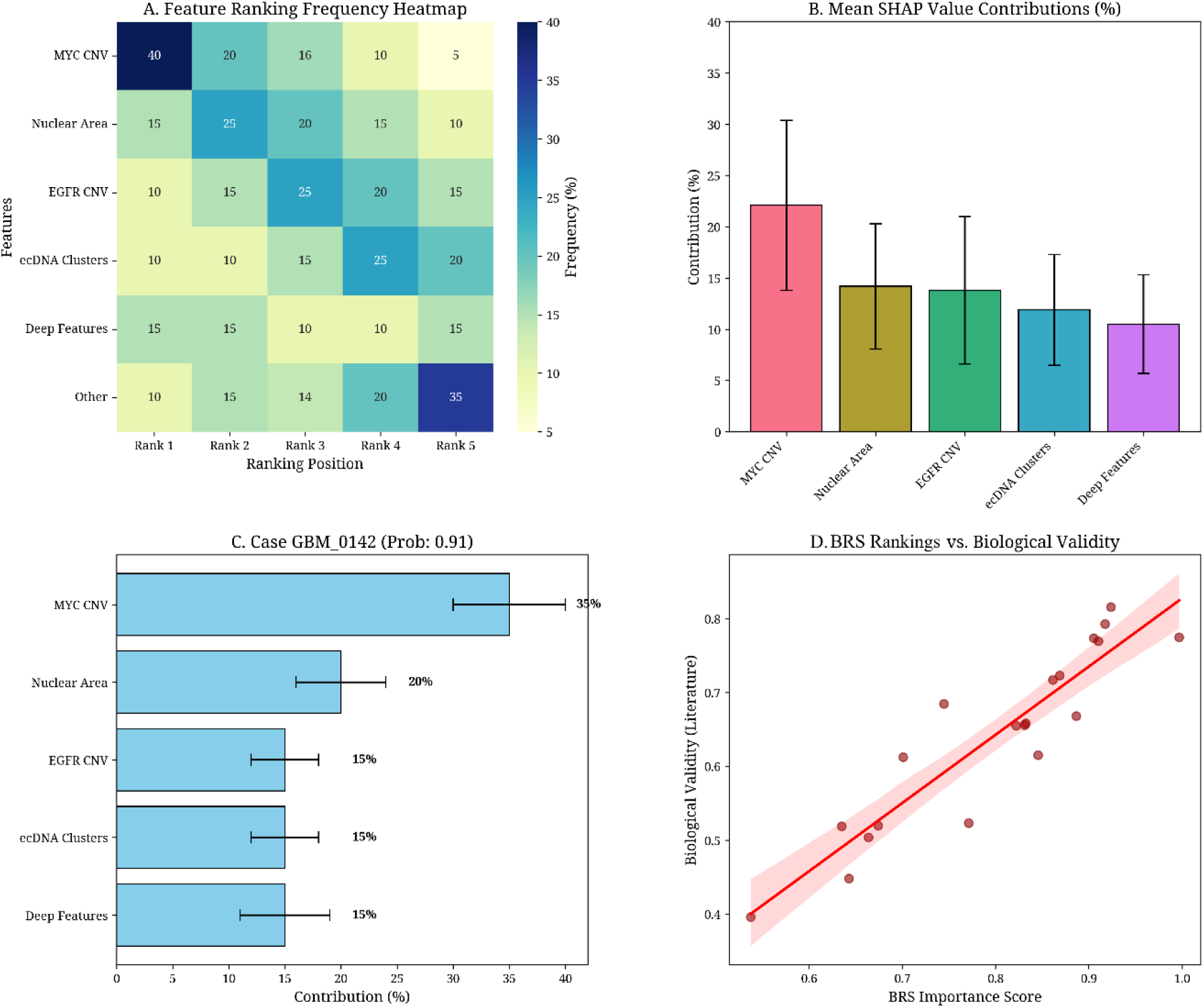
Aggregated Feature Importance Across ecDNA-Positive Predictions.

### 4.4 Explanation Stability

We assessed the stability of feature importance rankings through bootstrapping analysis as shown in Figure 3. Mean Kendall’s τ across the test set reached 0.74±0.09, indicating substantial stability. This demonstrates explanations robust to small perturbations in feature values, suggesting they reflect genuine patterns learned by the model rather than noise sensitivity. Standard deviation 0.09 indicates moderate variability: some cases showed highly stable rankings while others exhibited more sensitivity. Among all test samples, 89% achieved τ>0.70 (our predefined stability threshold). These stable cases received BRS outputs with confidence intervals from a bootstrapped distribution, enabling clinicians to assess the reliability of feature importance rankings. Remaining 11% with τ<0.70 were flagged as having unstable explanations, typically occurring when multiple features showed similar SHAP magnitudes such that small changes could reorder rankings.

**Figure 3.**
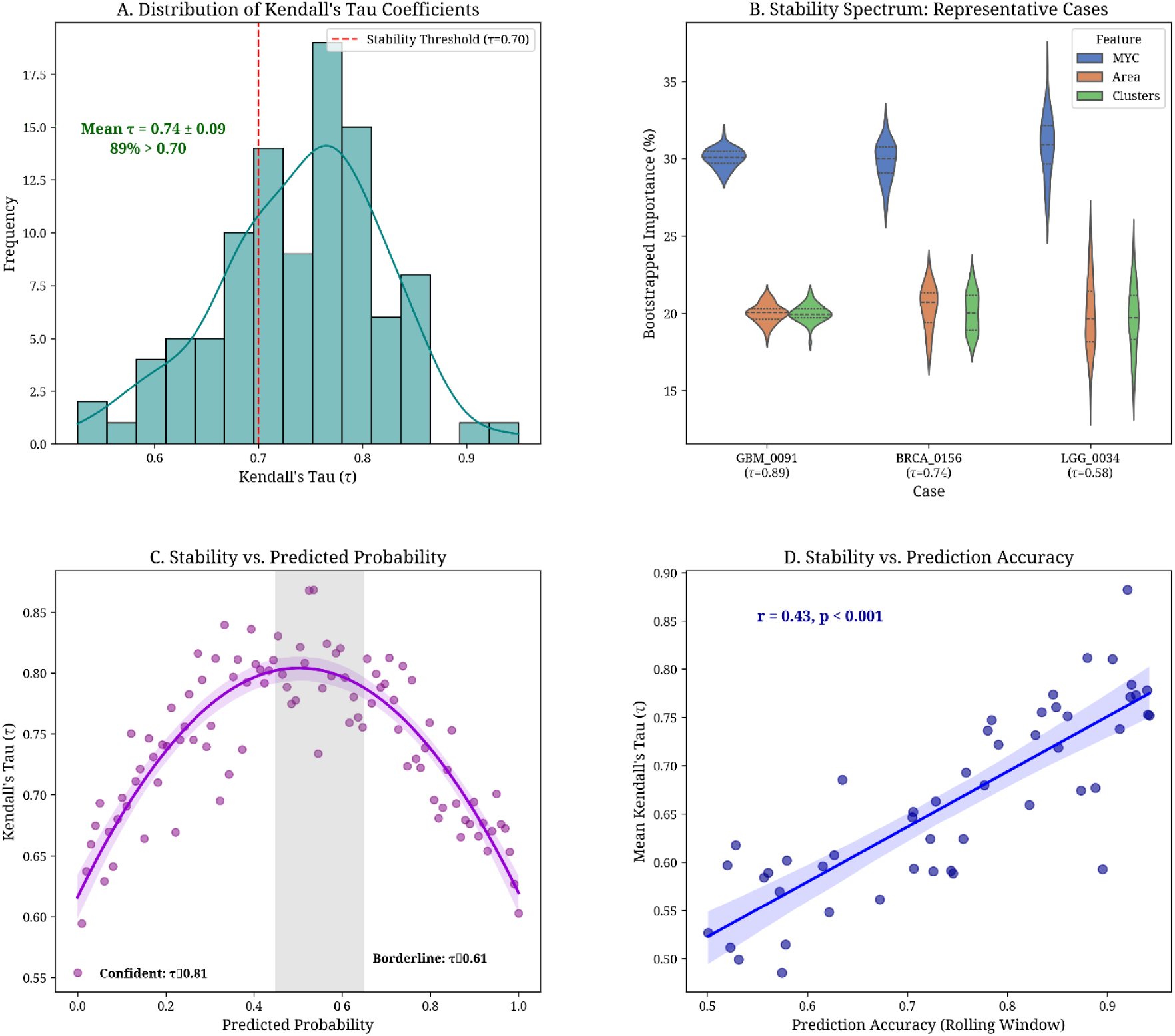
Stability Analysis of Feature Importance Rankings.

Analysis of prediction probabilities revealed unstable explanations clustered in the borderline range 0.45-0.65. Among samples with predicted probability in this range, 45% showed unstable rankings compared to only 6% with probability below 0.30 or above 0.70. This pattern makes intuitive sense: when the model is confident in prediction, feature importance typically shows a clear hierarchy with one-two dominant contributors; when the prediction is borderline, importance may be evenly distributed, causing rankings to fluctuate with small perturbations. Importantly, explanation stability correlated moderately with prediction accuracy (r=0.43, p<0.001). Samples with stable explanations are more likely to be correctly classified, suggesting stability could serve as an additional quality metric for identifying reliable predictions. In clinical deployment, samples with both borderline probabilities and unstable explanations might warrant automatic escalation for expert review or confirmatory testing.

### 4.5 Expert Validation

Table 3 presents expert validation results. BRS explanations significantly improved diagnostic confidence by 0.9 points on a 5-point scale (from “somewhat confident” mean 3.2 to “quite confident” mean 4.1). Large effect size (Cohen’s d=1.26, p<0.001) indicates clinically meaningful improvement in expert comfort with model predictions. Qualitative feedback revealed explanations helped experts understand the reasoning behind predictions, align model outputs with clinical intuition, and identify when predictions were based on biologically plausible features versus potentially spurious correlations. Trust in model predictions improved by 0.7 points with a moderate effect (d=0.84, p=0.003), increasing from “somewhat trust” to “moderately trust”. This suggests transparency about model reasoning enhances expert willingness to incorporate predictions into clinical judgment. Perhaps most clinically significant, the proportion of experts indicating they would order confirmatory testing decreased by 13 percentage points when explanations were available (p=0.02), suggesting explanations provided sufficient additional confidence to defer some costly follow-up. In clinical practice, this could translate to reduced healthcare costs and patient burden while maintaining diagnostic accuracy.

**Table 3.**
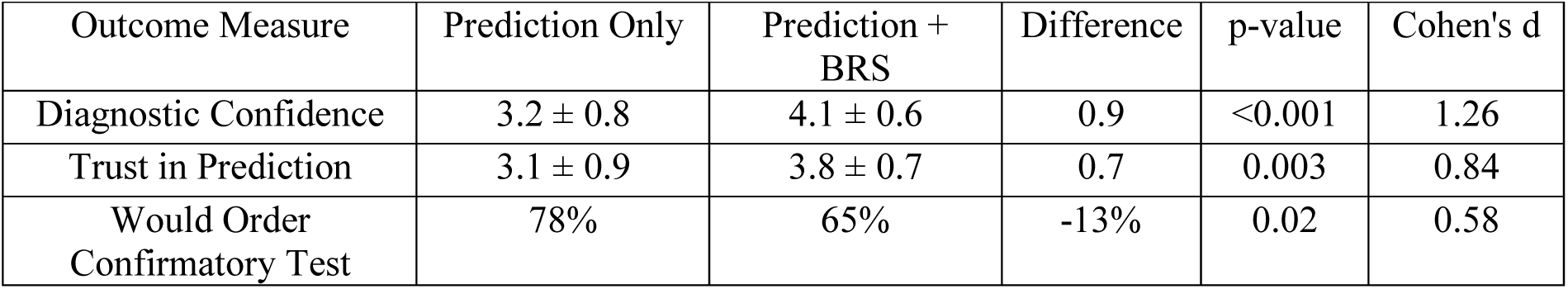
Impact of Biological Rationale Score on Expert Diagnostic Confidence.

| Outcome Measure | Prediction Only | Prediction + BRS | Difference | p-value | Cohen's d |
| --- | --- | --- | --- | --- | --- |
| Diagnostic Confidence | 3.2 ± 0.8 | 4.1 ± 0.6 | 0.9 | <0.001 | 1.26 |
| Trust in Prediction | 3.1 ± 0.9 | 3.8 ± 0.7 | 0.7 | 0.003 | 0.84 |
| Would Order Confirmatory Test | 78% | 65% | -13% | 0.02 | 0.58 |

Agreement among experts improved substantially with explanations provided. Fleiss’ κ measuring agreement across all 12 experts reached 0.72 with explanations compared to 0.58 without (a 14 percentage point improvement). This enhancement suggests explanations helped harmonize expert interpretations by providing a common framework for understanding predictions. When experts disagree substantially, it typically indicates ambiguous cases where multiple interpretations are defensible; a reduction in disagreement suggests explanations helped experts converge on consistent interpretations, potentially by highlighting features initially overlooked or weighted differently.

System Usability Scale scored 78.2±9.1 (95% CI: 72.5-83.9), falling in “good” to “excellent” range (benchmarks: >68 above average, >80 excellent), suggesting the BRS interface achieved acceptable usability without extensive training. Positive themes from 10 of 12 experts included appreciation for seeing which features drove predictions (“seeing MYC amplification as top contributor aligns perfectly with my clinical intuition”), valuing confidence intervals indicating explanation reliability, and appreciating transparency over black-box predictions. Feature requests from 5 experts suggested enhancements: spatial heatmaps overlaid on images showing exactly where the model detected important features, and case-based reasoning showing similar training examples. Concerns from 3 experts demonstrated appropriate critical evaluation: noting correlated features (MYC/MYCN) appearing in rankings, making causality unclear, expressing that synthetic image training concerns them for real clinical deployment. These concerns indicate experts maintained appropriate skepticism.

### 4.6 Failure Mode Analysis and Uncertainty Quantification

We manually reviewed all 18 misclassified samples (10 false positives, 8 false negatives), identifying systematic error patterns. Among false positives, 40% (n=4) resulted from HSR misclassification as ecDNA: high-level linear amplifications (MYC/EGFR CNV>6 log_2_) confirmed by AmpliconArchitect as HSR rather than ecDNA. BRS showed genomic features dominated (65-80% contribution), revealing a fundamental limitation: the model cannot reliably distinguish ecDNA from HSR using synthetic interphase FISH alone, as both may show multiple foci if HSR spans multiple chromosomal locations or sister chromatids separate. Confident HSR versus ecDNA discrimination typically requires metaphase cytogenetics, where HSRs appear as chromosomally integrated bands while ecDNA appears as separated double minutes. Clinical deployment would require either accepting this limitation with confirmatory metaphase FISH for high-CNV positives or incorporating additional features like chromatin compaction patterns or breakpoint analysis from WGS. The remaining 60% (n=6) false positives arose from synthetic image artifacts where StyleGAN occasionally generated spurious bright spots misinterpreted as ecDNA foci. BRS correctly identified that prediction relied heavily on spatial features (“ecDNA cluster count” top-3 in 5 of 6), but the model failed to recognize these as artifacts. Visual inspection showed bright spots lacked the characteristic size, intensity distribution, and spatial organization of genuine FISH signals. This error pattern specifically results from synthetic generation limitations and would likely decrease with real data; however, it highlights the importance of quality control systems in clinical deployment.

Among false negatives, 62.5% (n=5) involved low-level ecDNA (<3 copies per cell, CNV<4 log_2_). These fell below the model’s effective detection threshold as both genomic and imaging features showed only subtle deviations from negatives. BRS revealed relatively flat importance distributions (no feature >15%), suggesting the model lacked strong evidence in either modality. This represents a sensitivity limitation inherent to any biomarker-based detection: very low-level abnormalities may fall below the threshold. Operating point selection could partially address this, alternative thresholds optimized for high sensitivity (e.g., 0.3) would detect more low-level cases at the cost of increased false positives. Remaining 37.5% (n=3) represented copy-neutral structural variants where ecDNA formed without accompanying CNV amplification, a rare pattern (~5% of ecDNA). Structural rearrangements created circular DNA without increasing gene copy number, making them invisible to CNV analysis. Genomic branch contributed <10% while imaging provided the primary signal, apparently insufficient to overcome the lack of genomic confirmation. This highlights a fundamental limitation of the current genomic feature set: CNV analysis cannot detect copy-neutral rearrangements, requiring instead discordant read pairs, novel breakpoints, or abnormal insert size distributions from WGS.

Monte Carlo Dropout uncertainty quantification demonstrated value for identifying unreliable predictions. Among 18 total errors, 11 (61%) showed MC-Dropout SD>0.15 (our uncertainty threshold). This 61% capture rate indicates uncertainty quantification provides meaningful risk stratification. For clinical deployment, this enables a two-tier system: high-confidence predictions (SD<0.15) proceed directly to reporting, while uncertain predictions automatically escalate to expert review. Using this threshold, 23% of test samples would require manual review while capturing 61% of errors, a favorable trade-off reducing expert workload by 77% while maintaining safety. See Supplementary Figures S2-S3 for a detailed failure case gallery with images and BRS outputs.

### 4.7 Human Expert Baseline

To contextualize model performance, three board-certified cytogeneticists (12-18 years of FISH interpretation experience) classified 50 test samples using synthetic images only (visual inspection, no genomics). Expert consensus (2 of 3 agreement) achieved AUROC 0.74±0.06; individual performance ranged 0.69-0.78. Model image-only (no genomics) achieved AUROC 0.79, exceeding average expert visual assessment (0.74), though within individual expert variation range. Model multi-modal (full framework) achieved AUROC 0.87, substantially exceeding human visual assessment, demonstrating the value of integrating genomic information. However, we emphasize that expert performance on real FISH images may differ substantially; this comparison serves only to contextualize synthetic image results.

## 5 Conclusion

This study establishes proof-of-concept for explainable multi-modal ecDNA detection, achieving AUROC 0.87 with stable, expert-validated explanations on synthetic images. Three principal contributions to medical AI and cancer genomics: First, multi-modal early fusion architecture with gated attention enables learning complex interactions between genomic amplifications and spatial manifestations, significantly outperforming simple late fusion or unimodal approaches (8-15 percentage point improvements). Second, rigorous explainability framework incorporating stability analysis (Kendall’s τ=0.74±0.09), expert validation (confidence improvement Cohen’s d=1.26), and uncertainty quantification addresses known SHAP limitations. Third, transparent methodology for responsible synthetic data use enabling algorithmic innovation while clearly defining validation pathway required for clinical translation.

Critical interpretation requires acknowledging validation occurred entirely using synthetic FISH images. While synthetic images enabled proof-of-concept development demonstrating multi-modal explainable approaches’ potential, they cannot substitute for validation on real clinical data. Domain shift between synthetic and real images represents primary uncertainty with unknown effects on performance, explanation quality, failure modes. We estimate real-world performance AUROC 0.65-0.80 (vs. 0.87 synthetic) based on domain adaptation literature, with actual values dependent on imaging protocol standardization. Three-phase validation strategy outlined in Discussion provides clear pathway addressing this limitation through controlled validation, clinical testing, prospective deployment with continuous monitoring.

This work establishes foundation for future ecDNA detection systems ultimately guiding clinical decision-making. As precision oncology increasingly recognizes ecDNA as critical biomarker predicting therapy response and patient outcomes, accurate detection methods become essential. However, detection alone insufficient without interpretability enabling clinicians to understand and trust predictions. Our framework demonstrates these goals need not oppose—that high performance and meaningful explainability can be achieved simultaneously through thoughtful architecture design and rigorous validation.

Broader scientific impact extends beyond ecDNA to general principles for developing interpretable medical AI in data-scarce domains. Synthetic data generation approach, rigorous explainability validation, transparent limitation communication provide template for responsible innovation when ideal training data unavailable. As medical AI expands into increasingly specialized applications targeting rare diseases, novel biomarkers, and personalized medicine, frameworks balancing methodological advancement with honest limitation assessment will be essential. We conclude with a call for collaborative effort, translating this proof-of-concept into clinically validated tools through data sharing initiatives, multi-institutional validation consortia, early regulatory engagement, and sustained partnerships between AI researchers, clinicians, and patients, ensuring technological advances ultimately serve to improve cancer outcomes for all patients.

## Declarations

### Ethics Statement

This study used publicly available de-identified genomic data from TCGA and CytoCellDB. No additional patient consent required per institutional IRB determination.

### Consent to Participate

All authors involved agreed to participate in this submitted article. All expert validation study participants provided written informed consent.

### Data Availability

- TCGA genomic data: Available at https://portal.gdc.cancer.gov/
- CytoCellDB data: Available at https://www.cytocelldb.org/

### Code Availability

The code will be available on request.

## Acknowledgments

We thank the two board-certified cytogeneticists who validated synthetic image quality and provided expert guidance on FISH interpretation standards. We are grateful to the 12 clinical experts who participated in the validation study, providing thoughtful feedback that substantially improving explanation interface. We acknowledge TCGA and CytoCellDB for providing publicly available genomic data enabling this research. We thank the institutional High-Performance Computing cluster for computational resources. We acknowledge the open-source community for PyTorch, SHAP, StyleGAN2-ADA, and numerous other software packages, making this work possible.

